# Beyond library size: a field guide to NGS normalization

**DOI:** 10.1101/006403

**Authors:** Jelena Aleksic, Sarah Carl, Michaela Frye

## Abstract

**Background:** Next generation sequencing (NGS) is a widely used technology in both basic research and clinical settings and it will continue to have a major impact on biomedical sciences. However, the use of incorrect normalization methods can lead to systematic biases and spurious results, making the selection of an appropriate normalization strategy a crucial and often overlooked part of NGS analysis.

**Results:** We present a basic introduction to the currently available normalization methods for differential expression and ChIP-seq applications, along with best use recommendations for different experimental techniques and datasets.

We demonstrate that the choice of normalization technique can have a significant impact on the number of genes called as differentially expressed in an RNA-seq experiment or peaks called in a ChIP-seq experiment.

**Conclusions:** The choice of the most adequate normalization method depends on both the distribution of signal in the dataset and the intended downstream applications. Depending on the design and purpose of the study, appropriate bias correction should also be considered.

## Background

Normalization is a crucial step in any NGS data analysis pipeline. The purpose of this step is to transform the data in such a way as to make the data from different replicates and experimental conditions directly comparable. The transformation should ideally take into account any major factors that would shift or distort the data distribution, such as library size and differences in sample pools, as well as experimental biases introduced at any stage during the original experiment, library preparation or sequencing.

High-throughput genomics normalization methods were initially developed from approaches applied to the analysis of microarray data, where a number of systematic biases made it essential to perform correct data normalization. For example, in two-channel microarrays, it is known that the different dyes are incorporated with different efficiencies [1]. It therefore became essential to incorporate this into the experimental design by performing dye swaps. Dye bias also needed to be accounted for in the normalization process, by performing background correction, data transformation and between-array normalization [2]. In this way data were adjusted for systematic effects arising from the experimental biases rather than genuine biological differences, before proceeding to downstream analysis.

Some of the systematic biases inherent to microarray analysis are not relevant to NGS technology: For example, dye incorporation and background corrections are no longer issues. However, while it has been suggested that sophisticated normalization of datasets is no longer required for applications such as RNA-seq [3], further studies argue that the reality is not that simple, and that normalization still remains an important consideration [4, 5]. However, this is just one of a number of factors to consider, as different parts of the analysis pipeline such as alignment and quantification all impact the end results [6].

At a basic level, there is a need to normalise NGS data in order to account for different library sizes. Furthermore, systematic global differences in measurements may still be present in the samples, due to technical variation in the molecular biology leading up to the library preparation, and these must be corrected. Other biases are also apparent: for example, longer mRNAs are sampled more frequently, and in the case of ChIP, different sequences may be preferentially enriched or sequenced. Another complication is that for differential expression or enrichment analyses, any normalization method needs to remove the differences due to technical variation and sequencing depth, but not normalize away the genuine biological differences. This becomes difficult when the sequences of interest are a small subset of the total dataset, or when there are significant differences in expression between the different experimental conditions.

Choice of normalization method depends on the nature of the data being analysed, and different choices are therefore appropriate for different techniques, experimental conditions and data profiles. Importantly, different normalization methods can give very different results in terms of differentially expressed genes in the context of RNA-seq, or peak calling in the context of ChIP-seq. Here, we present an overview of the currently used NGS normalization methods, alongside guidelines for best use in different contexts.

## Results and Discussion

### Library size based methods

A number of NGS normalization methods take the size of each sequencing library as the basis for their normalization models. The reason for this is that even replicates from the same experiment sequenced together are usually sequenced to slightly different depths due to technical variability, and it is not uncommon to have very different sequence coverage between different experiments. Consequently some form of sequence read count transformation to make the library sizes comparable is a necessary step in NGS analysis.

#### Total count

The principle of total count (TC) based methods is that the total library count for each sample is used to calculate a normalization factor [7]. This method accounts exclusively for the differences in library size, and no other sources of variability. In its most basic form, if one library had 10 million reads, one would divide it by 10 to get a measure of reads per million (RPM). Alternatively, gene counts or binned read counts are divided by the number of total mapped reads in the sample, then multiplied by the mean total count across all the samples in the dataset. These have an equivalent effect – making the library sizes across samples comparable (Figure 1A and B).

**Figure 1.**
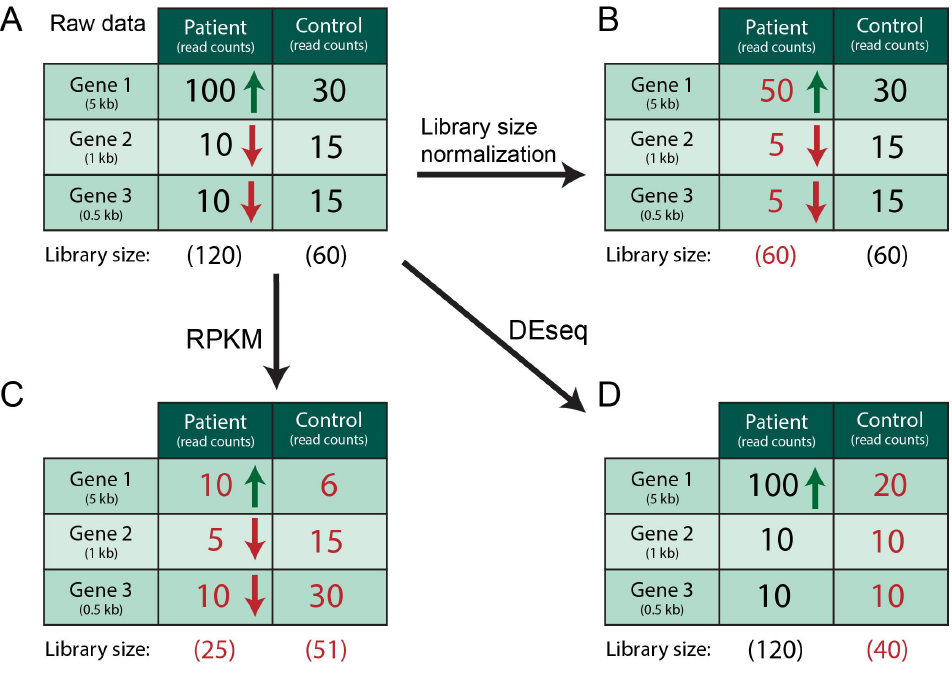
The conceptual basis of different normalization methods. For RNA-seq, starting from a raw data count (**A**), a number of different normalization methods can be employed. The commonly used library size (**B**), RPKM (**C**) and DESeq (**D**) methods are shown here. (**B**) For the library size method, the library sizes are equalized between the samples. (**C**) For the RPKM method, in addition to being library normalized, the sample counts are also divided by the gene length in order to account for gene length bias. (**D**) DESeq finds the majority of genes that do not appear to change expression (in this case genes 2 and 3), and uses these to calculate a scaling factor to equalise the libraries accordingly. The red and green arrows indicate the gene expression change direction indicated by the counts.

#### Variations of total count methods

There are a number of similar approaches that use counts of subsets of the library, rather than the total count, to calculate normalization factors. The upper quartile (UQ) method calculates a normalization factor based on the upper quartile of counts at mRNAs with non-zero counts. The method uses the sum of all upper quartile mRNAs in each replicate, and divides the replicates by this value [5]. The rationale behind the UQ approach is to focus on the higher and more stable signal, ignoring the variable low signal parts of the dataset. However, this assumes a similar distribution of counts between datasets being normalized. The median (Med) method is another simple variation, which calculates the median mRNA count for each replicate, and divides by that. The advantage is that it removes the potential influence of outliers.

#### Limitations of library size methods for RNA-seq

The library size based approaches make intuitive sense, since a library that is twice as large is expected to have approximately twice as many sequenced reads across each mRNA. This is often a valid assumption, particularly for the comparison of relatively similar experimental conditions or for normalizing between biological replicates of the same condition. Furthermore, one of the advantages of library size based methods is that they make no assumptions about the data: they do not require, for example, a certain proportion of mRNAs to not be differentially expressed.

However, problems arise when datasets with major differences in sample composition are compared. Total count-based normalization often biases the outcome when it is used for samples with large differences in expressed mRNA populations. If a particular set of mRNAs is very highly expressed in one experimental condition, but not the other, normalizing by a library size factor makes it appear that non-differentially expressed genes are in fact consistently downregulated, while minimising the genuine differences between the more highly expressed genes. As previously observed [4], this is often the case in RNA-seq, where identifying large differences in gene expression between experimental conditions is often the purpose of the experiment. Studying different sub-populations of genes is also known to produce this type of error. For example, focusing on tRNA genes within an RNA-seq experiment can pose problems because the proportion of tRNAs with respect to the total RNA composition may vary between experimental conditions. One way of adjusting for this is to use library size normalization on only a subset of the data; for example, normalizing by the total tRNA count, rather than the total library count.

While using this approach does remove some of the issues of library-based normalization, the issue of highly expressed genes still remains a problem, as these may still be present within sample sub-populations. Furthermore, since it is unknown in advance which genes will be highly differentially expressed between experimental conditions, it is not possible to remove this bias. It is instead generally preferable to use alternative normalization methods where possible.

#### Limitations of library size methods for ChIP-seq

Even libraries with a high depth of coverage have a finite number of reads; those reads are necessarily sampled from the population of molecules available in the sample. In a ChIP-seq experiment, the ChIP sample is expected to contain an enriched pool of DNA fragments that are bound by the protein of interest. This enrichment depletes the number of reads that will be sampled from other areas of the genome. When normalizing a ChIP library together with an input control library, which is expected to represent the distribution of background noise, scaling by total library size can inflate the noise in background regions, causing an artificially high FDR if a sample-swapping approach is used to estimate the empirical FDR [8, 9]. In both RNA-seq and ChIP-seq experiments, normalizing to total library size can lead to weak but biologically significant enriched regions or differentially expressed genes being missed.

### Scaling factors

Although the strategy of normalizing by total library size or reads per million is intuitively straightforward, as described above, problems can arise when the libraries being normalized for comparison have very different read distributions. Scaling factor based methods were developed to take this problem into account.

The general approach of scaling factor methods is to identify a subset of the signal that does not change between the experiments being compared and use that for normalization. For example, in the context of RNA-seq, it is assumed that the majority of mRNAs in a sample will not be differentially expressed. This population of stable mRNAs is used to normalize the rest of the samples, thus removing the issue of highly expressed genes. For ChIP-seq, most scaling factor approaches focus on identifying the background component of the ChIP dataset and scaling the input to match the background.

#### Scaling Factors in RNA-seq

For differential expression analysis of RNA-seq data, Robinson et al. suggest the trimmed mean of M-values (TMM) normalization method that uses the raw data to estimate appropriate scaling factors [4]. This can then be used in downstream analysis to account for the sampling properties of the data. This involves estimating relative abundances of different groups of genes and scaling the datasets accordingly. For example, if a group of liver-specific genes is highly expressed in RNA-seq data from liver, normalizing by library size alone would suggest that general housekeeping genes are significantly downregulated compared to, for example, the kidney. This is exactly what Robinson et al. observe, with 8% of housekeeping genes upregulated and 70% downregulated in the liver using standard normalization methods. They show that applying their scaling method (implemented in the edgeR Bioconductor package) reduces this bias, and makes the proportions more even between the samples. When the scaling normalization method is applied, 26% are upregulated and 41% are downregulated.

Another popular and widely used Bioconductor package, DESeq [10], and its more recent update DESeq2 [11], also use a variant of scaling factor normalization, based on the assumption that most genes are not differentially expressed. A scaling factor is computed for each sample, as the median of each gene’s scaling ratio (Figure 1D). The mRNA ratio is calculated as its read count over its geometric mean across all samples. The idea is that since most genes are not differentially expressed, they should therefore have a ratio of 1. If this is found not to be the case, a scaling factor is applied in order to make the data fit this assumption. This eliminates the problem of a set of highly expressed genes biasing the sequencing pool, since these are not taken into account during the normalization. However, the strategy is obviously only applicable if the assumption that most genes are not differentially expressed holds. The same constraint applies to the TMM normalization method, and is a general feature of scaling factor based methods, since they make use of invariant genes to scale the library accurately.

#### Implementations for ChIP-seq

A general solution to the scaling problem in ChIP-seq datasets involves separating the data into a signal component, corresponding to putative binding regions, and a background or noise component. A scaling factor is then calculated between the reads that make up the background of the ChIP and input libraries, the latter being assumed to be only background. This factor is then used to scale the entire dataset (Figure 2). The key parameter in this approach is how the background component is estimated, since the enriched regions are generally not known a *priori* [8]. Since ChIP-seq reads can show biologically interesting enrichment in any type of DNA, rather than just within genes or exons as is the case with RNA-seq, in a ChIP-seq analysis the genome is often divided into equally spaced bins or windows and the number of reads overlapping each bin counted. A number of different methods and implementations have been developed to assign each bin to either the enriched signal or the background; these can be roughly divided into methods that use a pre-determined cut-off to separate background from signal and methods that estimate the most appropriate threshold directly from the data.

**Figure 2.**
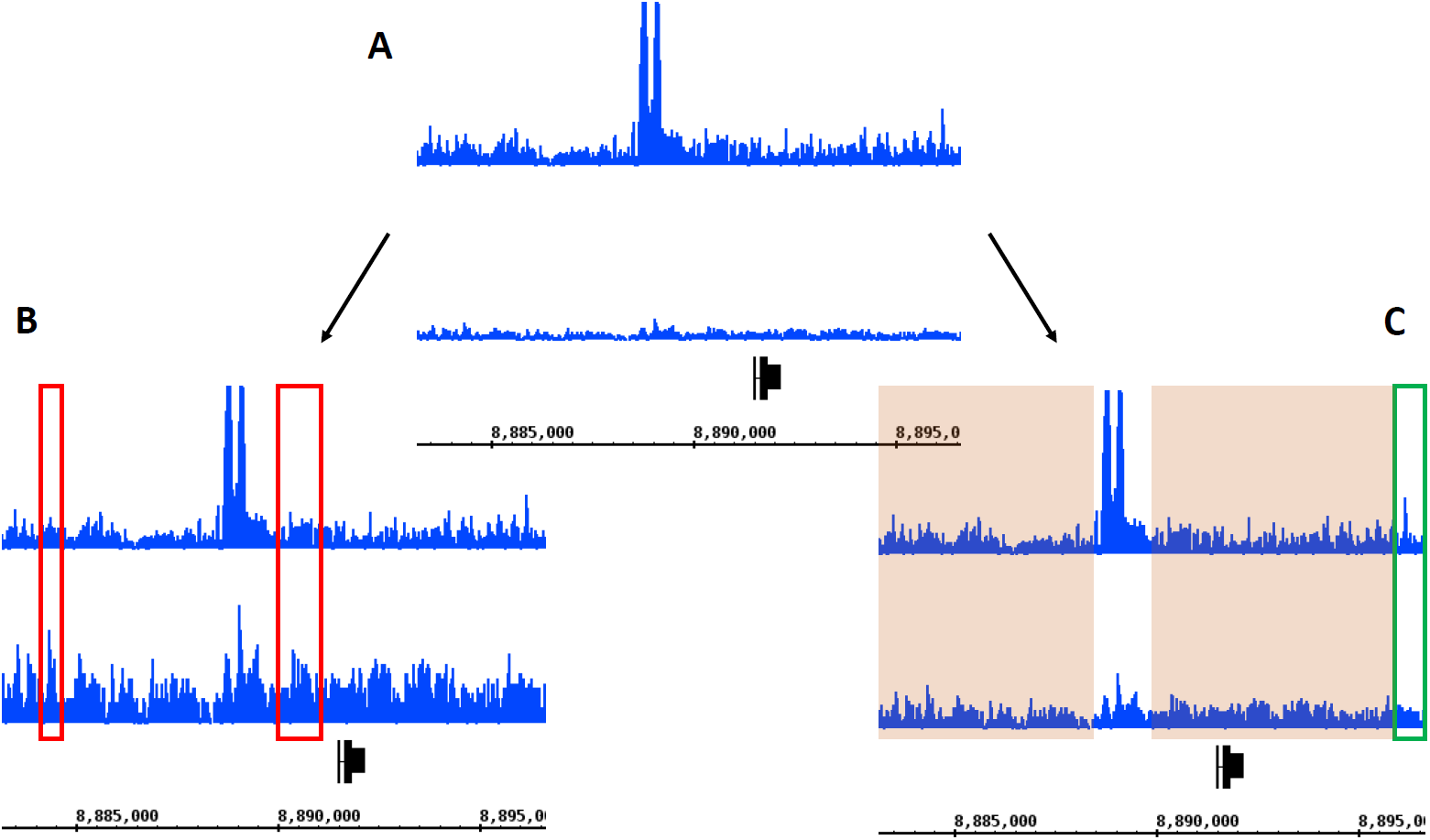
An illustration of the effect of total count scaling and scaling to the background in ChIP-seq. (**A**) A region of the Drosophila genome showing raw ChIP reads over input reads. In this dataset, there are roughly twice as many raw ChIP reads as input reads. (**B**) Scaling the input up to match the total size of the ChIP library results in amplification of noise in the input which could be interpreted as negative peaks, raising the FDR, as well as causing potentially real but weak peaks in the ChIP sample to be missed. Red boxes indicate areas of the input that could be called as negative peaks. (**C**) When the input is scaled to match the size of the background component of the ChIP signal, noisy regions of the input are less likely to be called as negative peaks and the sensitivity of peak-calling is improved. The green box indicates a peak that may be missed with total count scaling but that is detected when using this technique. The tan shaded areas show the background regions in each sample that are normalized to match.

Several early methods, which were integrated with the first ChIP-seq peak calling algorithms, fall into the first category. These include PeakSeq, CisGenome, and SPP, among others. The strategies for determining background reads with CisGenome and SPP are similar but complementary; both are based upon the observation that bins with enrichment in the ChIP sample will tend to have high read counts, while bins with a low number of reads will tend to fall into the background [8]. CisGenome designates all 100-bp bins with two or fewer reads as background and uses these bins to calculate the expected sampling ratio between sample and control in unbound regions. This ratio is then used in FDR estimation for each potential enriched bin [12]. SPP, on the other hand, estimates the background or nonspecific portion of the ChIP library by excluding highly enriched regions, defined as 1-kb regions with a number of tags that exceeds the expectation for a uniform Poisson distribution with a p-value less than 10^−5^ [13]. While these thresholding approaches are reasonable in most cases, it is not clear that they are appropriate for every dataset; indeed, the authors of CisGenome acknowledge that their methods may not be valid for datasets with low sequencing depth [12].

PeakSeq offers a more tuneable approach, using a simulated background Poisson distribution to identify potential enriched peaks as a first pass and then excluding a certain fraction of those peaks, *P_f_*, from the ChIP dataset to calculate the scaling factor between it and the input. While the *P_f_* parameter can be set independently for each analysis, providing more flexibility, it is unclear how to identify an optimal value. The authors demonstrate a significant difference in the results obtained when *P_f_* = 0 and when *P_f_* = 1, with 0 being the most conservative setting, yet they suggest that it should ideally be somewhere between these two extremes [14].

An alternative to specifying a fixed cutoff for determining the background component is to use properties of each dataset to estimate the optimal threshold directly. This adaptive approach recognizes that the selection of a count threshold for determining background bins implies a trade-off between bias and variance; smaller thresholds will tend to reduce bias more, as the resulting background bins will be very unlikely to contain weak but true regions of enrichment, but they will also display increased variance in the normalization factor estimates. Both NCIS and SES [8, 9] start by ranking bins by read count across the genome, then attempt to partition the bins into a subset containing only background and a subset in which enrichment is present. The normalization factor is then estimated at this partition point. NCIS searches for the minimum count threshold for which the ratio of ChIP counts in a bin to input counts in a bin is greater than that at the previous threshold and includes more than three-quarters of all bins. SES uses order statistics to search for the bin in which the maximum difference exists between the percentage of all input reads allocated to bins and the percentage of all ChIP reads allocated to bins.

A third approach, CCAT, models a ChIP library as a linear combination of signal and noise and estimates a noise rate from the input library, from which it is possible to calculate expected noise read counts in a given bin. Background bins are then defined as bins with read counts in the ChIP sample that are less than the expected noise counts [15]. However, there is evidence indicating that this algorithm may be sensitive to experimental artefacts, such as PCR overamplification during preparation of sequencing libraries, leading to significantly higher reads counts in the input library than in the ChIP library for some regions [8]. Comparisons suggest that these data-adaptive scaling factor methods can increase power and true positive detection when compared to both fixed-cutoff methods such as PeakSeq and methods that scale by total sequencing depth such as MACS [8, 9, 15].

### Bias Correction

As well as making library sizes comparable, ideally the normalization approach also considers any major biases arising from the experiment. These include biases based on gene length, as well as sequence content and accessibility. Here we present some popular methods for addressing these sources of bias.

#### Gene length biases and RPKM normalization

The Reads Per Kilobase per Million reads (RPKM) normalization was developed to specifically address transcript length bias and is a popular variant of library size based methods that is widely used for RNA-seq analysis (Figure 1C). It involves calculating a score relative to the size of each transcript, and is used as a way of facilitating a comparison of transcript levels while reducing the RNA-seq biases introduced by transcript length [16]. A similar concept, implemented in the Cufflinks software, is Fragments Per Kilobase of transcript per Million fragments mapped (FPKM). For single end data, RPKM and FPKM measures are the same, while in paired end data, FPKM treats paired reads as a single fragment [17].

The RPKM method is reported to introduce some variability, particularly in the case of genes expressed at low levels [7]. The basis of the problem is that longer transcripts have more of an opportunity to be randomly featured in an RNA-seq dataset, simply because of their length. Thus changes in long mRNAs may appear more significant, while changes in shorter mRNAs are more difficult to detect. The RPKM method involves dividing the library-normalised score by the total exon length of each transcript (e.g. dividing by 2 for a 2kb transcript). However, this causes issues for particularly small transcripts. For example, the counts for a 0.5kb transcript would be multiplied by 2, increasing the differences in the scores (Figure 1c). Since small transcripts are also likely to have low read counts, making them more prone to variability, this approach can result in the amplification of differences that are not genuinely biologically significant. Uneven transcript sequencing coverage is also a major issue, which can be unpredictable and vary significantly within and between samples, confounding analysis [18].

RPKM normalization can be helpful when the objective is to identify a globally representative set of differentially expressed genes. For example, gene length bias can have a big impact on gene ontology (GO) enrichment and network analysis, as lengths of different groups of functional genes are non-random. Which parts of the gene are relevant for normalization purposes also varies by experiment - for example, long introns can lead to a gene length bias in ChIP depending on how the peaks were assigned to genes, whereas it is transcript length that is relevant for RNA-seq. However, the downside is that it may remove genuine differentially expressed large genes in the process (an increase in false negatives), while at the same time adding false positive small genes whose originally small differences have as a result of normalization been flagged as significant.

#### Alternatives to RPKM

As mentioned, RPKM normalization introduces its own set of biases [7], and is considered not to be a valid between-sample normalization method [19]. However, the need to normalize for transcript length across samples still exists, and improved methods for achieving this have since been published. The Transcripts Per Million (TPM) measure is a modification of RPKM that is considered to be a more sound alternative for between-sample normalization [19, 20]. As well as taking into account the transcript length, it also considers the sequencing read length and the number of transcripts sampled, in order to estimate the relative molar concentration of RNA. TPM is an improvement, and is now included in a number of software packages such as RSEM, eXpress and sailfish.

In terms of differential expression, the BaySeq package uses Bayesian methods and includes gene length in the model [21]. This is important when comparing conditions within a species [4, 22], but really becomes essential in cross-species comparisons, where homologues may have different lengths and can therefore end up with very different read counts. DESeq2, the newer version of DESeq, also includes an option for calculating normalization factors for gene length, which can be used to correct for gene length bias in a differential expression analysis [11].

#### Sequence biases

A separate but related issue in normalizing ChIP-seq libraries, or indeed any type of genome-wide count data, is that of correcting for biases that may be introduced by local variations in sequence mappability, sequence composition or chromatin accessibility. In early ChIP-seq experiments, it was assumed that the background followed a uniform Poisson distribution; however, analysis of input chromatin libraries showed that the background tag distribution exhibited significantly more clustering than expected [13]. These “background peaks” can arise for a number of reasons, including the tendency of sequencing chemistry to favour sequences with higher GC content [23], differences in chromatin accessibility resulting in some regions being sheared more readily during sonication, and differences in mappability, as sequences that map to more than one region of the genome are often excluded from analysis. Since these sources of bias can be correlated with functional genomic features (for example, exons tend to have a high GC content), it is important to correct for them to avoid systematically under- or over-estimating the significance of peaks in certain regions of the genome [14, 24].

The most straightforward way to correct for potential biases is to use a matched input control prepared under the same conditions as the sample, since this should be subject to all of the same sources of error. Many normalization and peak-calling methods do just this, either dividing the sample reads by the scaled input reads or subtracting the input reads from the sample reads at each position [12–14, 25]. Others, including the popular peak-caller MACS, use a sample-swapping strategy to estimate the FDR by calling both sample peaks over the control and control peaks over the sample [26]. However, there is evidence to suggest that simply using an input sample to correct for local biases is not sufficient, as using different replicate inputs can give different results and even dividing one input library by another can yield regions of apparent enrichment [24]. Another possible control for a ChIP-seq experiment is a mock-IP, performed using an antibody that is not expected to bind to anything in the sample. While this approach may reduce false positives more efficiently than an input control, it can also be subject to bias and greater variability, as mock-IP samples often have a very low DNA yield and are thus vulnerable to PCR amplification artefacts [27].

A commonly proposed solution to this problem is to correct libraries using independently determined data such as mappability profiles and calculations of % GC content across the genome. PeakSeq provides code to calculate mappability for fragments of a user-specified size across any given genome, and the resulting maps are used in the peak-calling process [14]. Several other tools, including MOSAiCS, BEADS and ZINBA, also make use of this type of data. Both MOSAiCS and ZINBA utilize mixture modelling, in which additional information such as GC content and mappability are treated as covariates [28]. ZINBA allows the incorporation of other data, such as copy number variations. This approach does not separate normalization and peak-calling into two different steps, but rather uses all available covariates to model three components of the data: background, enriched regions, and zero-counts corresponding to regions with insufficient coverage or low mappability [29]. On the other hand, BEADS does not call peaks but focuses solely on bias correction. It transforms raw data by weighting reads by mappability and GC content, as well as calculating a local correction from input data; the algorithm is quite flexible and can also be applied to input datasets themselves [24]. However, in order to apply it to ChIP datasets, potential enriched regions must first be identified using another method, as the GC content is calculated based on the background only. The resulting corrected data can then be normalized by library size or using one of the scaling factor methods outlined above. Bias correction using mappability and GC-content can also be a suitable solution for situations when an input dataset is not available. Other biases include more frequently mapping alleles that match the reference genome better than those that include sequence variants - these can be simulated using a known sequence set, and should be taken into account for any applications where variant-specific expression is important.

### Normalization methods comparison

We applied some of the more popular normalization methods to published datasets from human and fly, to test their impact on ChIP-seq and RNA-seq results.

#### Impact of normalization on RNA-seq

In the case of RNA-seq, we compared three popular normalization methods: library size total count, DESeq scaling factors, and RPKM normalization. We selected representative publicly available datasets, including single cell profiling of human zygotes and oocytes [30], and a comparison of *Drosophila* adult males and females from the modENCODE project [31, 32]. It was hoped that the fly data in particular would give us a good indication of false positives and false negatives, since male-female differences are well characterised from a genomic perspective.

In terms of the differentially expressed genes, DESeq and library size normalization gave very similar results for both human and fly data (Figure 3A). While some minor differences were observed, over 90% of the identified genes were the same in both cases. The RPKM method on the other hand consistently identified a smaller number of differentially expressed genes with both human and fly data. The difference in the human data was more pronounced, with RPKM normalization identifying 1500 differentially expressed genes, compared to around 2300 total identified by both DESeq and library size normalization methods. Encouragingly, the great majority of RPKM-identified genes were also identified by the other two methods. This shared set of genes was also found to have similar expression patterns across all three experimental methods, meaning that at least in the case of these datasets, the library size normalization method did not lead to major data skew.

**Figure 3.**
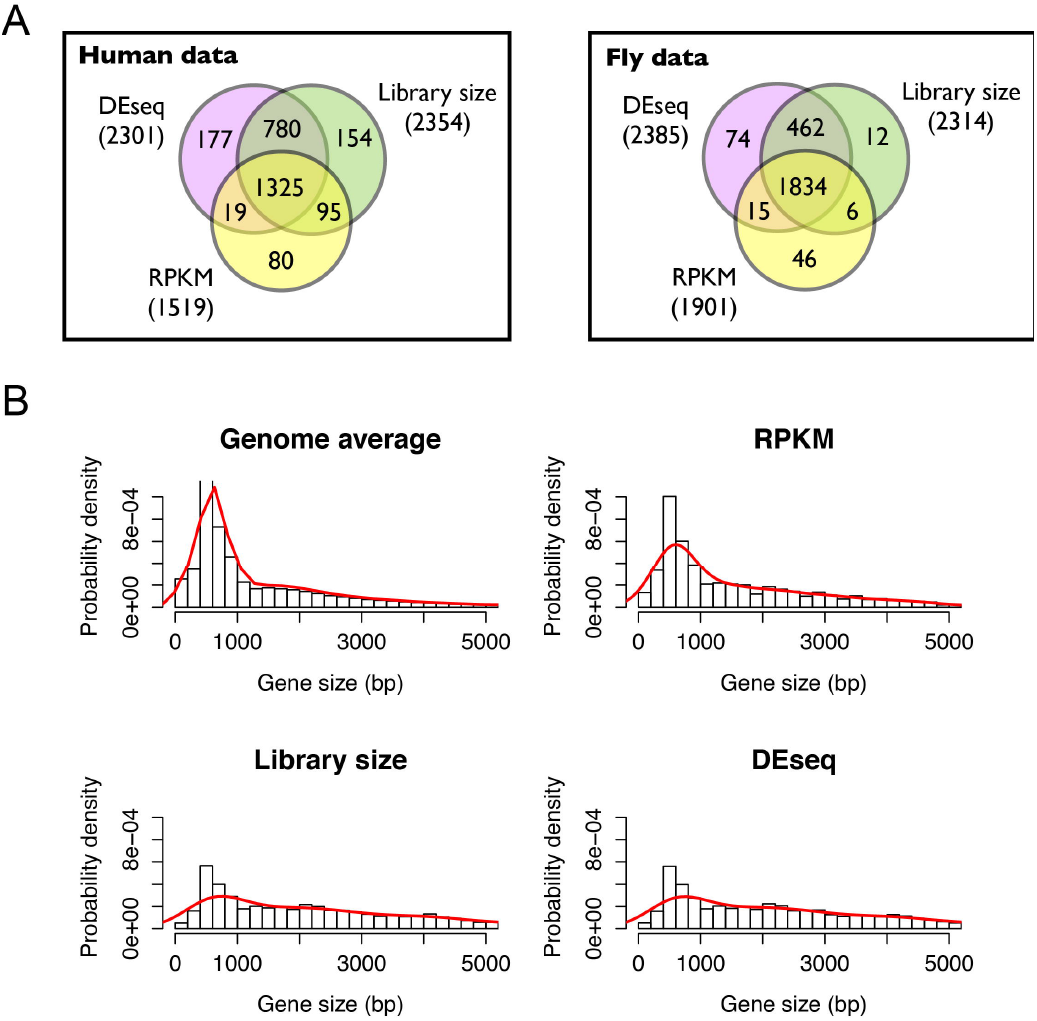
A comparison of different RNA-seq normalization methods. (**A**) Differentially expressed genes were called using the DESeq package, but with different normalization methods. RNA-seq data from a single cell oocyte and zygote study was used for human [30], and modENCODE data comparing adult males and females was used for fly [31]. While there is a large amount of overlap across all normalization methods, RPKM shows a much lower sensitivity than the other two methods. (**B**) The probability density of total exon sizes for different transcripts is shown for the genome as a whole, as well as for the differentially expressed genes identified using RPKM, library size and DESeq normalization methods. The RPKM normalization method more closely matches the gene size distribution in the whole genome, while library size and DESeq methods contain more long genes than would be expected at random.

Assessing the sizes of the differentially expressed genes found that the mean total exon length was significantly higher than the genome average for all three normalization methods, though more so for DESeq and library size. However, it is clear from the size distributions (Figure 3B) that the RPKM method more closely matches the genome distribution, while library size and DESeq are more skewed towards longer genes.

Which normalization method should be applied depends both on the exact dataset in question, and the application for which the data is being analysed. Since it is possible to introduce a large systematic skew if there is a group of highly expressed genes present within a particular experimental condition, it seems prudent to use DESeq scaling factors by default to avoid this. However, there are also applications where gene length bias will significantly skew the results, and this might become the more important consideration. For example, any systems biology applications that require a relatively non-biased sampling of genes, such as GO enrichment or network analysis, would benefit from normalization that takes into account gene length bias. On the other hand, if sensitivity is an issue, DESeq or library size will likely pick up a larger number of differentially expressed genes. Conversely, to identify a robust set of reliable genes for further biological validation, it may be worth trying out multiple methods and choosing the genes that are commonly identified.

#### Impact of normalization on ChIP-seq

We tested the effects of three normalization methods with both human and Drosophila ChIP-seq datasets: TC-based scaling, the SPP method, and the NCIS method. Sequencing reads were downloaded from a ChIP-seq experiment for the D. melanogaster embryonic segmentation transcription factor Knirps (Kni) and for the human liver-specific transcription factor CEBPA [32–34]. The Kni dataset consists of three ChIP replicates and three input replicates, while the CEPBA dataset consists of four ChIP replicates and four input replicates. In both cases, all ChIP and input replicates were pooled; while this does not allow for assessment of reproducibility, and while we would generally recommend analysing all replicates independently, it is the recommended procedure for using MACS, which we chose as a baseline peak-caller for evaluating different normalization methods. All reads were aligned against the most recent release of the respective reference genome.

In order to visualize the distribution of reads in each dataset, mapped reads were counted in bins across the genome (500 bp for human and 100 bp for *Drosophila*) and input bins were scaled by the ratio determined by each of the normalization methods tested. ChIP reads versus input reads were then plotted for each bin (Figure 4). From these plots, it is immediately clear that different ChIP-seq datasets can have very different signal distributions; in comparison to the Kni dataset, the CEBPA dataset contains more bins with higher counts in the ChIP sample than in the input, which could be considered to be the bins containing the ChIP enrichment signal. Both datasets also contain a number of bins with higher counts in the input sample than in the ChIP, these outlier bins with very high counts in the input sample could be due to PCR overamplification. Over each scatterplot, we plotted lines at y = x, which represent the points at which bins have the same number of reads in both the ChIP and the input samples. Simplistically, bins falling above this line could be considered as candidate peaks, since they have more counts in the ChIP sample than in the input. These plots illustrate the effect that different normalization methods have on determining the threshold for the ChIP signal, as well as the differences between datasets.

**Figure 4.**
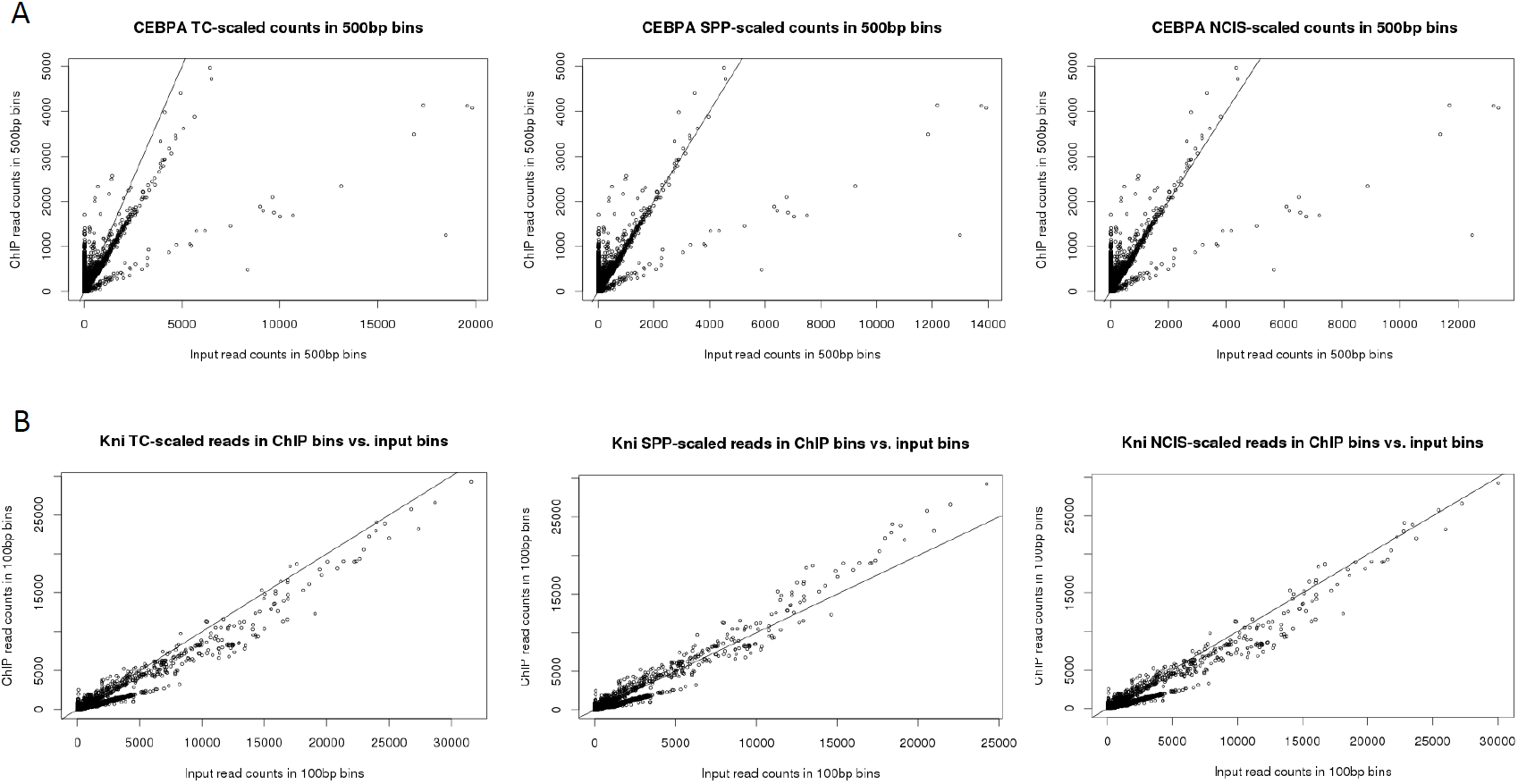
Binned ChIP counts versus input counts after normalization. (**A**) CEBPA ChIP data from human livers [34] in 500-bp bins. (**B**) Knirps modENCODE ChIP data from Drosophila embryos [33] in 100-bp bins. Both datasets were normalized by scaling binned input counts by the ratios derived from total library counts (TC), SPP and NCIS. A line at y = x is plotted on each graph to illustrate the point where bins have equal numbers of counts in the ChIP and input samples, roughly corresponding to the threshold of ChIP signal. The effect of different normalization strategies can be observed in the number of bins that fall above and below the line in each plot.

We wanted to decouple the effect of normalization from the peak-calling process, as many of the tools that implement different normalization methods also implement different algorithms for peak-calling. To do so, we used MACS, a popular ChIP-seq peak caller whose default setting is to scale the larger of the two datasets (either ChIP or input) to the smaller dataset using total library size. A patch available in MACS v1.4.4 (https://github.com/taoliu/MACS/tree/macs_v1) allows users to specify a custom scaling ratio with which to scale the control library. We first confirmed that the default scaling factor used by MACS matches the ratio between the ChIP and control libraries calculated, either by NCIS or simply by dividing the total number of reads in the ChIP library by the total number of reads in the input library, then ran MACS using the default p-value threshold of 1e-5 on each dataset to test the effect of TC normalization. We repeated the analysis using the --*ratio* option and specifying each alternative scaling ratio, with all other parameters remaining the same, to test the effect of each other normalization method.

For the CEBPA dataset, scaling by the total library size resulted in 110069 peaks and 3559 negative peaks. Each alternative scaling method both increased the number of positive peaks and significantly decreased the number of negative peaks. Since MACS calculates the FDR by performing a sample swap and determining the number of peaks called in both ChIP versus control and control versus ChIP at any given p-value [26], this represents both an increase in numbers of peaks and a decrease in the estimated FDRs. The NCIS method, which estimated the lowest scaling ratio, was the most effective at maximizing positive peaks and minimizing negative peaks. For this dataset, NCIS estimates that 67% of bins in the ChIP sample correspond to the background component, with 33% containing ChIP signal. The differences between the TC method and the other methods can be seen in Figure 4A; scaling by the total library size clearly results in an overestimation of the background, with many more bins falling below the diagonal, while the other scaling factors fit the shape of the data more appropriately. A summary of the parameters, ratios, and results for each analysis is presented in Table 1.

**Table 1.**
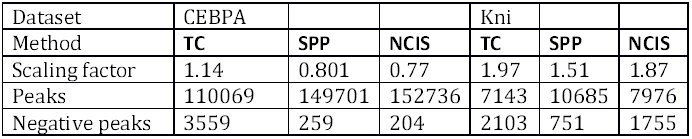
Scaling ratios derived from each normalization method for each ChIP-seq dataset, along with positive and negative peaks called by MACS when using each ratio.

For the Kni dataset, 7143 peaks and 2103 negative peaks were called using the TC normalization. Using the NCIS-estimated scaling factor resulted in a slight increase in positive peaks and decrease in negative peaks; however, the estimated FDRs were still quite high (∼20-30%) for the majority of peaks, reflecting the high number of apparently enriched regions in the input. The relatively modest improvement seen with the NCIS scaling factor may be explained by the fact that NCIS estimates that 95% of the 100-bp bins in the ChIP sample for this dataset correspond to the background component, meaning that in this case the difference between the background size and the total library size is small. Indeed, the NCIS scaling factor, 1.87, and the TC ratio, 1.97, are fairly close. This reflects what is seen in Figure 4B, where few bins fall above the diagonal. On the other hand, the SPP scaling factor estimate for this dataset is considerably lower, resulting in the greatest number of positive and lowest number of negative peaks. The different effects of these methods on the CEBPA and Kni datasets highlight the importance of plotting the data to understand its distribution and testing various methods of normalization before proceeding with downstream analysis.

## Conclusions

Normalization is an essential step in any NGS analysis, and one that significantly impacts the results of downstream analysis. In the context of RNA-seq, we generally recommend the use of scaling factors such as those implemented in DESeq and edgeR, as long as the data matches the required assumption that the majority of genes do not change expression. However, the optimal normalization method also depends on the application for which it is being used. When particular biases present a challenge for the downstream analysis, appropriate normalization methods should be used to counter those biases; for example, methods that take into account gene length bias. This becomes particularly crucial in cross-species analysis, where homologue length may change between species.

In the context of ChIP-seq, we find that TC-based scaling between a ChIP and matched input sample is often overly conservative and can artificially raise the FDR. We recommend NCIS as an easy-to-use tool to estimate the percentage of ChIP bins in the background component and the corrected scaling factor between the input and the background. However, as our analysis of two different datasets shows, the background estimation can vary widely depending on the distribution of the data. Consequently, it is useful to plot the data beforehand and to test several different tools. We emphasise that the majority of tools that include an estimation of the background for normalization also use their own algorithms for calling peaks, meaning that the final set of peaks will also vary based on, for example, whether SPP or MACS is used as a peak caller. Nonetheless, it is important for researchers to understand how their data is normalized by such tools and to adjust it if necessary; tools like MACS which allow for flexibility in specifying a scaling ratio can be very helpful in this regard.

For sensitive applications where data bias is likely to be a major issue, we recommend using an appropriate bias correction method, for example, correcting for transcript size in RNA-seq applications. However, we also emphasise that correct experimental design in both ChIP-seq and RNA-seq applications is essential, and will go a long way towards countering the biases inherent in the techniques.

## Methods

### RNA-seq normalization

The datasets used were from single cell profiling of zygotes and oocytes in human [30], and a comparison of *Drosophila* adult males and females from the modENCODE project [31, 32]. Raw sequencing reads were downloaded from GEO, including zygote and oocyte data from series GSE36552 for human, and datasets GSM451804 and GSM451805 for fly. The SRA files were decompressed using SRA toolkit 2.3.5. The sequencing reads were aligned to the genome using Tophat v2.0.9 allowing for 2 splice mismatches (-m 2) [35]. Human data was mapped to the human genome version GRCh37/hg19 with Ensembl release 71 gene annotations, while fly data was mapped to genome BDGP R5/dm3, Ensembl release 75 gene annotations [36–38]. Only uniquely mapping reads were used for the analysis. Sam files were created using SAMtools [39], and htseq-count was used to get summary counts for each transcript [40].

DESeq was then used to call differentially expressed transcripts with the same settings in all cases [10], with the only variable being the normalization method used. The scaling factor method is native to DESeq, so the analysis for this was run using software defaults. For the library size methods, the DESeq scaling factors were replaced with library sizes in millions of reads for each replicate. For the RPKM method, the scaling factors were replaced by library sizes, and the counts were adjusted for exon length of each transcript before reading them into DESeq. All differentially expressed genes were called at the same cut-off (p-value <= 0.05).

### ChIP-seq normalization

Raw sequencing reads for the *Drosophila* and human ChIP-seq experiments were downloaded from GEO [GEO:GSE23147] and ArrayExpress [ArrayExpress: E-TABM-722], respectively. All reads for replicates from the same experimental condition were pooled. Reads were mapped against the D. melanogaster April 2000 release (dm3) for *Drosophila* and the December 2013 (hg38) release for human, obtained from the UCSC Genome Browser (http://genome.ucsc.edu/) [36–38]. In the case of the human data, all unassembled chromosomes were excluded from further analysis. Bowtie2 was used for mapping with the default settings [41].

100-bp or 500-bp bins across the *Drosophila* and human genomes, respectively, were generated using the BEDTools makewindows utility [42]. Counts of reads in all bins were then calculated using the BEDTools coverage utility. Binned counts were plotted in R v3.1.0 using RStudio v0.98. NCIS and SPP were also run in R v3.1.0 using RStudio v0.98. Peaks were called using MACS v1.4.4 run with Python v2.7. For library-size based normalization, the following command was used: *macs –t treatment_file.bam –c control_file.bam –f BAM –g genome –n name -- keep-dup all --to-large*. For each other method, the appropriate calculated scaling ratio was used by substituting the --to-large option with the --ratio option.

## Authors’ Contributions

J.A. reviewed and performed the comparative analysis of RNA-seq normalization methods. S.C. reviewed and performed the comparative analysis of ChIP-seq normalization methods. J.A. and S.C. drafted the manuscript, with help from M.F. All authors read and approved the final manuscript.

## Acknowledgments

The authors would like to acknowledge the members of the CSCR/CSBC Bioinformatics Journal Club for insight and discussion during the preparation of this manuscript. Special thanks to Bettina Fischer for discussion, advice and help with R, and to Richard Smith-Unna for gene length normalization discussions. We would also like to thank Steve Russell for his insightful comments about the manuscript and help with preparing the draft. S.C. is supported by a Wellcome Trust 4-year Ph.D. Studentship in Developmental Biology and a Cambridge Overseas Trust Scholarship. J.A. is supported by European Research Council funding.

